# Failure to detect mutations in *U2AF1* due to changes in the GRCh38 reference sequence

**DOI:** 10.1101/2021.05.07.442430

**Authors:** Christopher A. Miller, Jason R. Walker, Travis L. Jensen, William F. Hooper, Robert S. Fulton, Jeffrey S. Painter, Mikkael A. Sekeres, Timothy J. Ley, David H Spencer, Johannes B. Goll, Matthew J. Walter

**Affiliations:** Division of Oncology, Department of Internal Medicine, Washington University School of Medicine, St. Louis, MO, USA; Siteman Cancer Center, Washington University School of Medicine, St. Louis, MO, USA; McDonnell Genome Institute, Washington University School of Medicine, St. Louis, MO, USA; The Emmes Company, 401 North Washington Street, Suite 700, Rockville, MD 20850, USA; Moffitt Cancer Center, Tampa, FL, USA; Division of Hematology, Department of Medicine, Sylvester Comprehensive Cancer Center, University of Miami School of Medicine, Miami, FL, USA; Department of Pathology and Immunology, Washington University School of Medicine, St. Louis, MO, USA

## Abstract

The U2AF1 gene is a core part of mRNA splicing machinery and frequently contains somatic mutations that contribute to oncogenesis in MDS, AML, and other cancers. A change introduced in the GRCh38 version of the human reference build prevents mutations in this gene from being detected by many variant calling pipelines. We describe the problem in detail and show that a modified GRCh38 reference build with unchanged coordinates can be used to ameliorate the issue. This reference is available at https://zenodo.org/record/4684553 (doi:10.5281/zenodo.4684553)

---

The *U2AF1* gene encodes a core component of the messenger RNA splicing machinery that is frequently mutated in MDS and other cancers^1–3^. Though predominantly associated with hematopoietic cancers (73%), mutations are also recurrent in lung tumors (6.5%) and have been reported in 24 other tumor types. Specifically, mutations at residues S34 and Q157 have been shown to promote exon skipping and are confirmed driver mutations contributing to cancer pathogenesis^4–7^.

As part of the National Myelodysplastic Syndrome (MDS) Natural History Study (NCT02775383), we used a targeted gene panel to sequence bone marrow samples from 120 patients either diagnosed with MDS or suspected to have MDS^8^. Of these patients, 38 were eventually confirmed to have MDS or MPN. Initial analyses looking at sequencing quality metrics revealed coverage levels and mutation frequencies that closely matched expectations, with one exception: mutations in the *U2AF1* spliceosome gene are typically observed in nearly 10% of MDS or Myeloproliferative Neoplasm (MPN) patients, but only 2 of the 38 (5.2%) MDS or MPN patients in this group had such mutations, both at the Q157 hotspot^1,9^. While this deviation was not significant, it was concerning that both mutations had far lower sequence coverage than would be expected, as did the rest of the *U2AF1* gene. Manual inspection with IGV revealed that this poor coverage extended to a 150 kilobase region of the genome where the majority of reads had a mapping quality of zero (Fig. 1)^10^. This region spans the *CSS*, *U2AF1, FRGCA*, and *CRYAA* genes.

**Figure 1:**
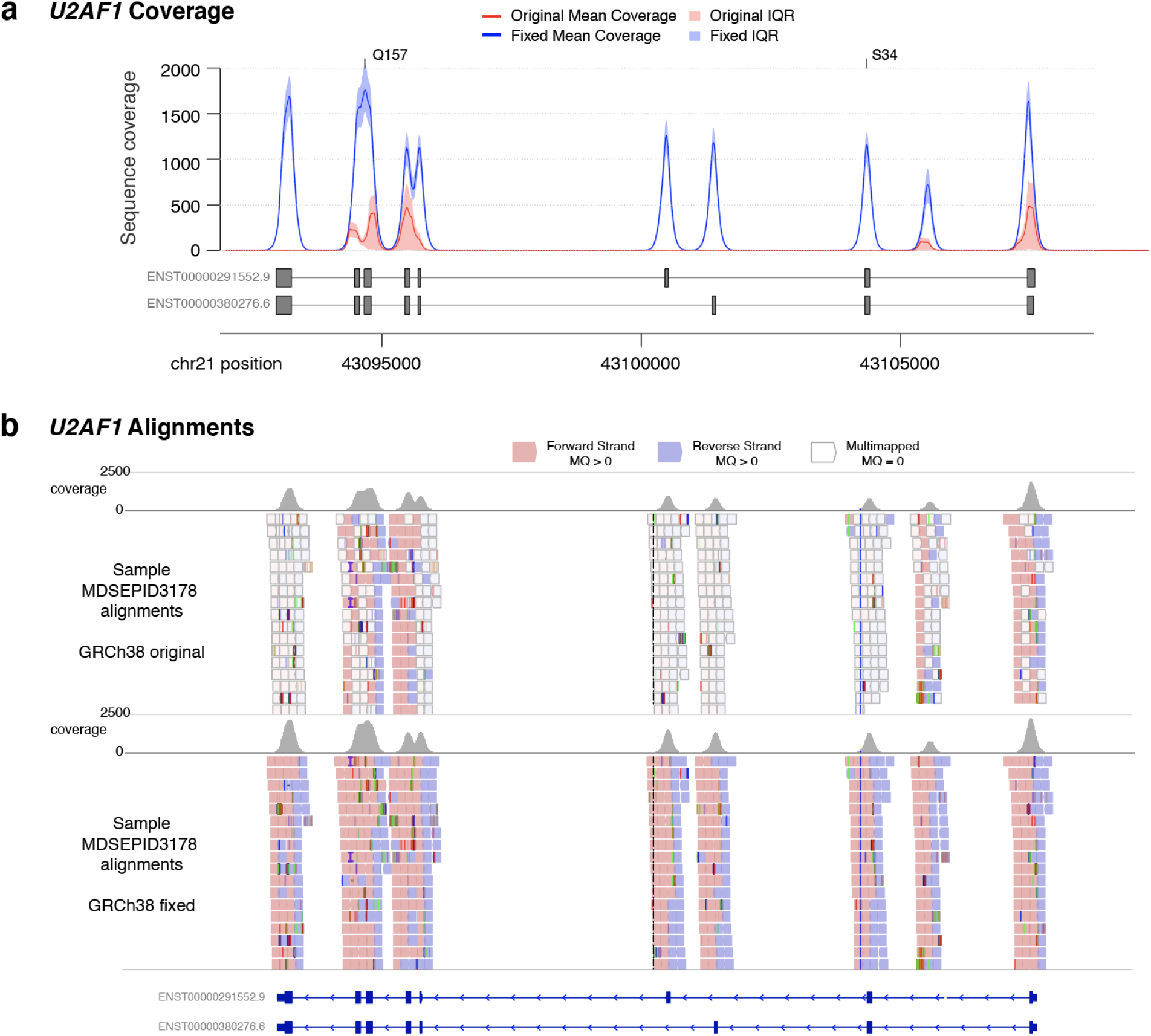
Alignment issues across the *U2AF1* locus. Panel a: Sequence coverage of reads with mapping quality greater than zero across the *U2AF1* gene for 120 bone marrow samples sequenced after capture with a custom reagent. The mean coverage for realignments to GRCh38 are shown in red, while the mean coverage for alignments to our modified GRCh38 reference is shown in blue. Shading indicates the interquartile range for each. Exons from the two primary protein-coding isoforms are shown below, and the locations of hotspot mutations at amino acids S34 and Q157 are indicated. Panel b: An IGV view of sequence reads, with alignments to GRCh38 at top and alignments to the modified reference at bottom. Grey bars at top show overall coverage. Reads colored in white indicate multimapped reads, with mapping qualities of 0, while red and blue colored reads have higher quality alignments.

Further investigation revealed that in the GRCh38 reference build, content added to the p-arm of chromosome 21 (chr21:6427259-6580181) contained sequence that nearly exactly replicated the sequence of the *U2AF1* locus (chr21:43035875-43187577). The same issue does not exist in prior reference builds GRCh36 or GRCh37. After consultation with members of the Genome Reference Consortium (GRC), it was determined that a BAC clone (https://www.ncbi.nlm.nih.gov/nuccore/FP236240.8) was incorrectly added to the reference genome, creating this duplicate sequence. This resulted in the alignment algorithm, BWA-MEM, splitting the reads among these two loci and assigning mapping quality scores of zero to reads that were multi-mapped^11^. This lowered the overall coverage substantially (Fig. 1, Supplementary Fig. 1), and as such alignments are typically excluded or down-weighted during variant calling, explained the paucity of mutations we observed in *U2AF1*, especially at the S34F position.

To address this issue, we created a modified version of GRCh38 that maintains the coordinate system, but replaces the new duplicate sequence on chromosome 21p with “N” characters. We realigned the data to this reference, and observed a substantial increase in coverage and mapping qualities across the affected region. Over exons of *U2AF1*, the coverage rose from a median of 0.3x to a median of 1,195x. This enabled the discovery of an additional *U2AF1* mutation (S34F) in this cohort (Supplementary Table 1).

To validate this finding in an orthogonal data set, we examined data from Acute Myeloid Leukemia (AML) patients sequenced for The Cancer Genome Atlas paper^12^. The data in this study were originally aligned to genome build GRCh36, and 6 mutations in the *U2AF1* gene were reported from exome sequencing. We downloaded exome sequence data that had been realigned (using BWA-MEM) by the core Genomic Data Commons pipelines against the GRCh38.d1.vd1 reference^11,13^. The GDC also provided somatic variant files from four different algorithms, none of which reported mutations in the *U2AF1* gene in any sample. Running our own somatic variant calling pipeline (https://git.io/JYlpq) on these alignments also did not reveal any *U2AF1* mutations.

Review of the alignments from these TCGA cases revealed the same coverage and mapping quality issues on chromosome 21. One sample lacked any *U2AF1* reads in the available exome data due to unrelated sequencing problems (the TCGA consortium only identified the mutation using orthogonal assays). After realigning the data from the remaining 5 cases to our modified version of GRCh38, the same somatic pipeline as above identified all 5 evaluable mutations in *U2AF1* (Supplemental Table 1).

The reference genome is essential to modern cancer genomics, but problems with these assemblies have the potential to cause both false positives and negatives. In this study, we describe the latter, where changes introduced in the GRCh38 human reference build cause mutations in a key driver gene to be missed with standard analysis approaches. GRCh38 was first released in late 2013, but widespread adoption lagged somewhat, so some studies involving myeloid malignancies (TCGA, Beat AML) have been spared this issue because they used older versions of the reference. Nonetheless, we expect that large cancer databases, including the NCI’s Genomic Data Commons, may be missing many *U2AF1* mutations due to this artifact. This also has clinical implications, because *U2AF1* mutations have strong associations with prognosis and clinical trials of splice-modulating drugs are being planned or are underway^5,14,15^.

We have reported these findings to the Genome Reference Consortium (https://www.ncbi.nlm.nih.gov/grc/human/issues/HG-2544) but as there are no patches or new releases currently scheduled for the human reference genome, the problem remains unresolved in the current release of the GRCh38 reference (GRCh38.p13). In order to ameliorate the problem, we have generated a corrected reference that is coordinate-compatible, and the differences in the resulting alignments are almost entirely in the masked and *U2AF1* regions. This means it can be used interchangeably with the standard GRCh38 reference in applications where detection of *U2AF1* mutations is critical, including sequencing of hematological cancers or studies of spliceosome dysfunction. The reference fasta file is available at https://zenodo.org/record/4684553 (doi:10.5281/zenodo.4684553).

## Data Availability

Sequence data from the MDS cohort is available in dbGaP under accession id phs###### (submission in progress). TCGA AML data is available via the Genomic Data Commons at https://portal.gdc.cancer.gov/

## Code Availability

Code used for variant calling is available at https://github.com/genome/analysis-workflows. Links in the Supplementary Methods are to the specific commits of the relevant workflows used.

## Acknowledgements

Supported by a National Cancer Institute (NCI) Research Specialist Award (R50 CA211782, to Dr. Miller), a Genomics of Acute Myeloid Leukemia Program Project grant (P01 CA101937, to Dr. Ley), and the Edward P. Evans Foundation (to Dr. Walter). The National MDS Natural History Study has been supported by US Federal Government Contracts HHSN268201400003I and HHSN268201400002I from the National Heart, Lung, and Blood Institute and additional funding by the National Cancer Institute to the participating member clinical centers in the NCORP and NCTN. This work has been supported in part by the Tissue Core Facility at Moffitt Cancer, an NCI designated Comprehensive Cancer Center (P30-CA076292). We thank Nancy DiFronzo for leadership of the National MDS Natural History Study. The National MDS Natural History Study thanks the study participants, as well as the investigator teams at the participating clinical sites.

## Author Contributions

Conceptualization: CAM, JRW, TLJ, WFH, RSF, DHS, JBG, MJW. Sample acquisition was led by JSP. Data analysis was performed by CAM, JRW, TLJ, WFH. Visualization: CAM, JRW, TLJ. Supervision and Funding: TJL, JBG, MAS, MJW. Draft writing and editing: CAM, JBG, TLJ, MJW. All authors read and approved the final manuscript.

---

**Supplementary Figure 1:**
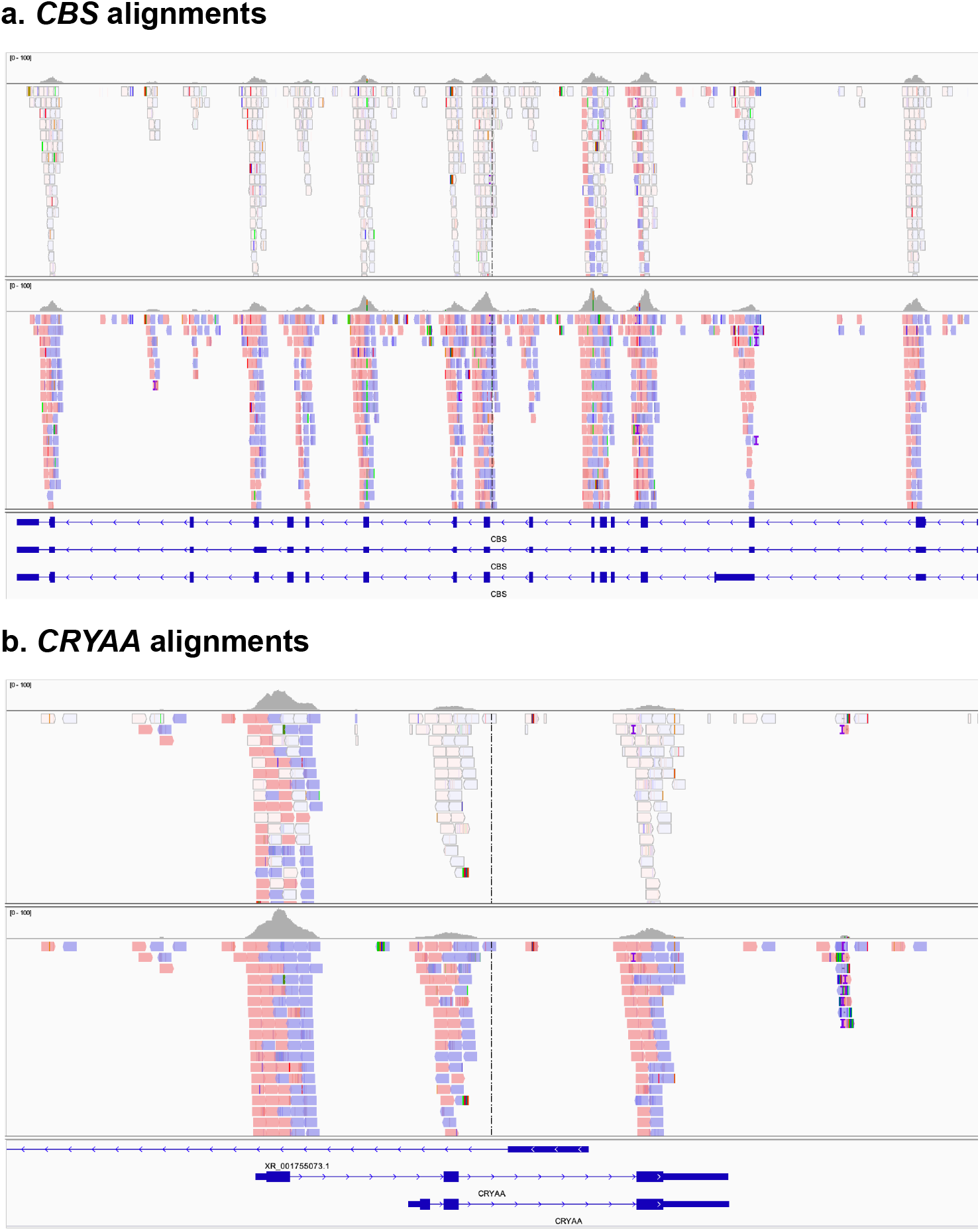
Examples of poor coverage of other genes in the affected region. Example shown is from exome sequencing of sample TCGA-AB-2912-11A. IGV views show sequence reads, with alignments to GRCh38 at top and alignments to the modified reference at bottom. Grey bars at top show overall coverage. Reads colored in white indicate multimapped reads, with mapping qualities of 0, while red and blue colored reads have higher quality alignments. Panel a: reads aligning to the *CBS* gene. Panel b: reads aligning to the *CRYAA* gene. (*FRGCA*, the other gene in the affected region, was not targeted by this exome reagent)

## Supplemental Information

### Methods

Sequencing reads were realigned to our modified reference using BWA-MEM, followed by sorting and deduplication, as detailed in a CWL workflow at https://git.io/JYbGl. Variants in the MDS cohort were called in single-sample mode using VarScan2 with params “--min-coverage 20 --min-reads2 5 --min-var-freq 0.05 --strand-filter 1”. Somatic variants in the AML cohort were called using an ensemble approach in the somatic pipeline described in detail at https://git.io/JYbGM

